# A Video Segmentation Pipeline for Assessing changes in Pupil Response to Light After Cannabis Consumption

**DOI:** 10.1101/2023.03.17.533144

**Authors:** Benjamin Steinhart, Ashley Brooks-Russell, Michael J. Kosnett, Prem S. Subramanian, Julia Wrobel

## Abstract

Due to long-standing federal restrictions on cannabis-related research, the implications of cannabis legalization on traffic and occupational safety are understudied. Accordingly, there is a need for objective and validated measures of acute cannabis impairment that may be applied in public safety and occupational settings. Pupillary response to light may offer an avenue for detection that outperforms typical sobriety tests and THC concentrations. We developed a video processing and analysis pipeline that extracts pupil sizes during a light stimulus test administered with goggles utilizing infrared videography. The analysis compared pupil size trajectories in response to a light for those with occasional, daily, and no cannabis use before and after smoking. Pupils were segmented using a combination of image pre-processing techniques and segmentation algorithms which were validated using manually segmented data and found to achieve 99% precision and 94% F-score. Features extracted from the pupil size trajectories captured pupil constriction and rebound dilation and were analyzed using generalized estimating equations. We find that acute cannabis use results in less pupil constriction and slower pupil rebound dilation in the light stimulus test.

## 1 Introduction

Cannabis consumption has been steadily increasing in the United States and consequentially, accidents involving cannabis have increased. The percentage of fatalities from motor vehicle accidents involving cannabis or a combination of cannabis and alcohol have doubled from 2000 to 2018 (Lira et al., 2021). Despite increased availability of cannabis, reliable methods for detection of recent cannabis use have been elusive. The Standard Field Sobriety Test has been used to reliably detect impairment due to alcohol since 1981 but shows limited ability to detect cannabis consumption (Papafotiou et al., 2005; Downey et al., 2012). Additionally, for those with daily use of cannabis, tetrahydrocannabinol (THC) can remain in the blood days or even weeks after cannabis consumption contributing to its unreliability as a biomarker of recent cannabis use. When examining risk of crash, THC was not a statistically significant predictor (Brubacher et al., 2019). Identifying a reliable, objective biomarker of recent cannabis use has proven challenging, but pupillary response may offer an avenue for detection that outperforms typical sobriety tests and THC concentrations.

The pupillary light reflex is a physiological constriction of the pupil that occurs in proportion to light stimulation of the retina. When light hits the retina, the signal is carried by the optic nerves to the midbrain, where equal neural impulses are generated and sent to both pupillary sphincter muscles to cause pupillary constriction. Various stimuli (e.g., pain or arousal) and drugs (e.g., those that alter sympathetic stimuli to iris dilator muscles or parasympathetic output from pupilloconstrictor neurons) may influence the size of the pupil and the extent of the pupillary light reflex. Reports have inconsistently characterized the impact of cannabis smoking or vaping and THC (tetrahydrocannabinol), the primary psychoactive agent in cannabis, on pupil diameter or the pupillary light reflex. Depending on the investigation, acute cannabis smoking or vaping has been associated with a decrease (Brown et al., 1977; Ortiz-Peregrina et al., 2020), increase (Stark et al., 2003; Merzouki et al., 2008; Shahidi Zandi et al., 2021) or no change (Fant et al., 1998; Newmeyer et al., 2017) in static pupil diameter. The pupillary light reflex, as assessed by the relative degree of illumination-induced pupil constriction, was decreased after acute cannabis consumption in some studies that utilized quantitative infrared pupillometry (Fant et al., 1998; Campobasso et al., 2020).

Two retrospective analyses of qualitative eye examinations conducted on apprehended drivers by law enforcement officers associated the presence of a positive blood test for THC with an increased prevalence of “dilated pupils” and “rebound dilation” (non-sustained light-induced pupil constriction) based on subjective assessments made without ocular instruments (Logan et al., 2016; Hartman et al., 2016). Drug Recognition Expert protocols utilized by many law enforcement jurisdictions rely on subjective estimation of pupil size and rebound dilation to support an assessment that an individual is under the influence of cannabis (IACP, 2022). However, the assessments by Drug Recognition Experts have a subjective element, and the conditions under which they are administered may vary. A standardized test of pupil changes could improve the ability of law enforcement professionals to assess driving-related cannabis use by replacing their observations with precise, objective, and replicable measurements of pupil changes.

The objective of the present study was to determine if data on pupil size and the pupillary light reflex extracted from infrared videography could be used to determine the effect of acute cannabis smoking in occasional and daily cannabis use. In order to quantify pupillary light reflex it was necessary to segment the videos of pupils under light stimulus. We developed a novel multi-step video processing procedure to ensure accurate segmentation of pupils from the infrared videography. Several challenges were encountered in the segmentation of videos produced by the stand-mounted goggles. Namely, pixel intensities from eyelashes, image edges, and irises can match intensities of pupils from the same image. Additionally, pupil locations and size vary based on goggle headset fit. Lastly, there are calibration points (bright white dots) that can obscure the pupil (Supplement Figure 1). To obtain standard results across participants, we developed a video segmentation pipeline that utilizes image processing and statistical techniques to obtain smooth pupil size trajectories over the duration of the light response test. From these smooth trajectories we calculate interpretable summaries of pupil change and compare them across cannabis use groups.

## 2 Methods

### 2.1 Data Collection

Healthy adults aged 25-45 were recruited to our Colorado data collection site between October 2018 and February 2020. Participants were enrolled into one of three groups: (1) daily cannabis use defined as smoking or vaping cannabis flower product at least one time per day, every day of the week for 30 days prior to enrollment (*N* = 33); (2) occasional cannabis use defined as smoking or vaping cannabis flower product on at least one day but no more than two days per week in the 30 days prior to enrollment (*N* = 36); and (3) no use defined as having not used cannabis in the month prior to enrollment (*N* = 32). All subjects appeared for initial data collection after a required interval of at least 8 hours of abstinence from cannabis that was verified by a multi-day subject consumption diary. See Brooks-Russell et al. (2021) for additional screening and enrollment details.

The study used a within-subject observational design with two assessment periods denoted “pre” and “post”. For subjects in the occasional or daily use group, “pre” and “post” assessments were completed before and after smoking cannabis, respectively. These subjects were observed to smoke or vaporize cannabis flower during a 15-minute interval and were instructed to smoke adlibitum “the amount you most commonly use for the effect you most commonly desire.” Subjects in the no use group were invited to relax for the equivalent amount of time. At both the pre and post assessment periods blood draws, a set of pupillary light response tests administered using goggles with infrared videography, and other data not reported here was collected. The post-use measurement of the pupillary light response occurred a mean of 71.36 (SD:4.56) minutes after the start of the ad-libitum smoking interval.

#### 2.1.1 Pupil Video Data

Pupil data was collected using goggles and protocol developed by Ocular Data Systems, Inc (Pasadena, CA). Ocular Data Systems designed their SafetyScreen™ goggles with infrared videography aimed to replicate, in a controlled setting, a set of light response tests similar to those conducted by Drug Recognition Experts. Videos of participants’ eyes during the light response test were collected twice, once at the pre and once at the post event.

Subjects sat upright with their face positioned in contact with the SafetyScreen™ goggles mounted on a stand. Infrared video of pupils were recorded from cameras inside the goggles. The procedure was conducted in a dimly lit room to reduce outside light contamination. Subjects were instructed to stare directly into the screen inside the goggles before and during illumination of their eyes with high intensity visible light emitted within the goggles for the duration of the test. Twenty-second videos of the eye with frame rates of 30 frames per second were recorded to capture pupil behavior under the light stimulus. The exact duration of the light test varied across individuals (median [IQR] = 14.4 seconds [12.7-15.0]). Similarly, the exact start and end time of the light test for each subject was not recorded in the SafetyScreen™ software and was estimated post-segmentation; our procedure for estimating the test start and end times is described in Section **2.2.4**.

To obtain a ground truth against which to select segmentation parameters and validate results, we manually segmented pupils from 30 frames of 60 randomly selected videos. Right and left eyes were manually segmented in equal proportions. The images were spread out in equal time intervals capturing the pupil at different points in the light stimulus test. Manual segmentation was performed using Cell Profiler (Lamprecht et al., 2007), a graphical user interface commonly used for segmentation of biological data (Supplement Figure 2). Manually segmented images were divided into a 70/30 percent split for tuning/validating segmentation parameters. Sixty-six images were removed due to lack of pupil visibility that occurred while subjects were blinking (Supplement Figure 1**D**) giving total sample sizes of 1217 and 517 manually segmented images in the tuning and validating sets, respectively.

### 2.2 Segmentation-Analysis Pipeline

The subsections discussed below explain a combination of image processing and statistical techniques that were used to segment pupil data. Our combination of established techniques allow for customization to solve different segmentation problems that can be described using ellipses. Each step could be swapped out and improved upon without modifying downstream analysis.

#### 2.2.1 Notation and Summary

We use the notation *I*_*itfe*_(*p*) to represent the grayscale intensity for a single 450×400 pixel image at pixel *p*. The subscript *i* ∈ {1, 2, … 101} represents the subject, *t* ∈ {0, 1} identifies the pre (*t* = 0) or post (*t* = 1) assessment timepoint, *f* ∈ {1, 2, … 600} represents the frame number, and *e* ∈ {0, 1} indicates the left (*e* = 0) or right (*e* = 1) eye. Pupil segmentation produces a binary mask for each image, which we denote 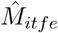 for masks estimated via our automated pipeline. We denote manually segmented masks 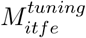 or 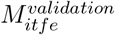 to represent manually segmented masks used tuning segmentation parameters or validation of the segmentation pipeline, respectively.

Our segmentation pipeline produces an estimated pupil mask 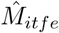 at each frame of each video, from which we calculate pupil major axis size, denoted *Y*_*itfe*_. The result is that for every video, there are estimates for pupil size at every frame in the video for each eye, which creates a pupil size trajectory. We fit a median additive regression model to remove trajectory outliers, which returns smooth estimates for major axis size, 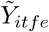. Then, from each smooth pupil size trajectory, we extract interpretable single-number summary features described in Section **2.2.5** and then compare these features across cannabis use groups.

#### 2.2.2 Segmentation

To segment pupils, each image was smoothed, intensity transformed, binarized using Otsu’s multi-level thresholding algorithm, and then fit with an ellipse to produce a binary mask. This process is depicted stepwise in Figure 2 and described in detail below. Each step Figure 2 modifies the same raw image shown in Figure 2**A**.

**Figure 1:**
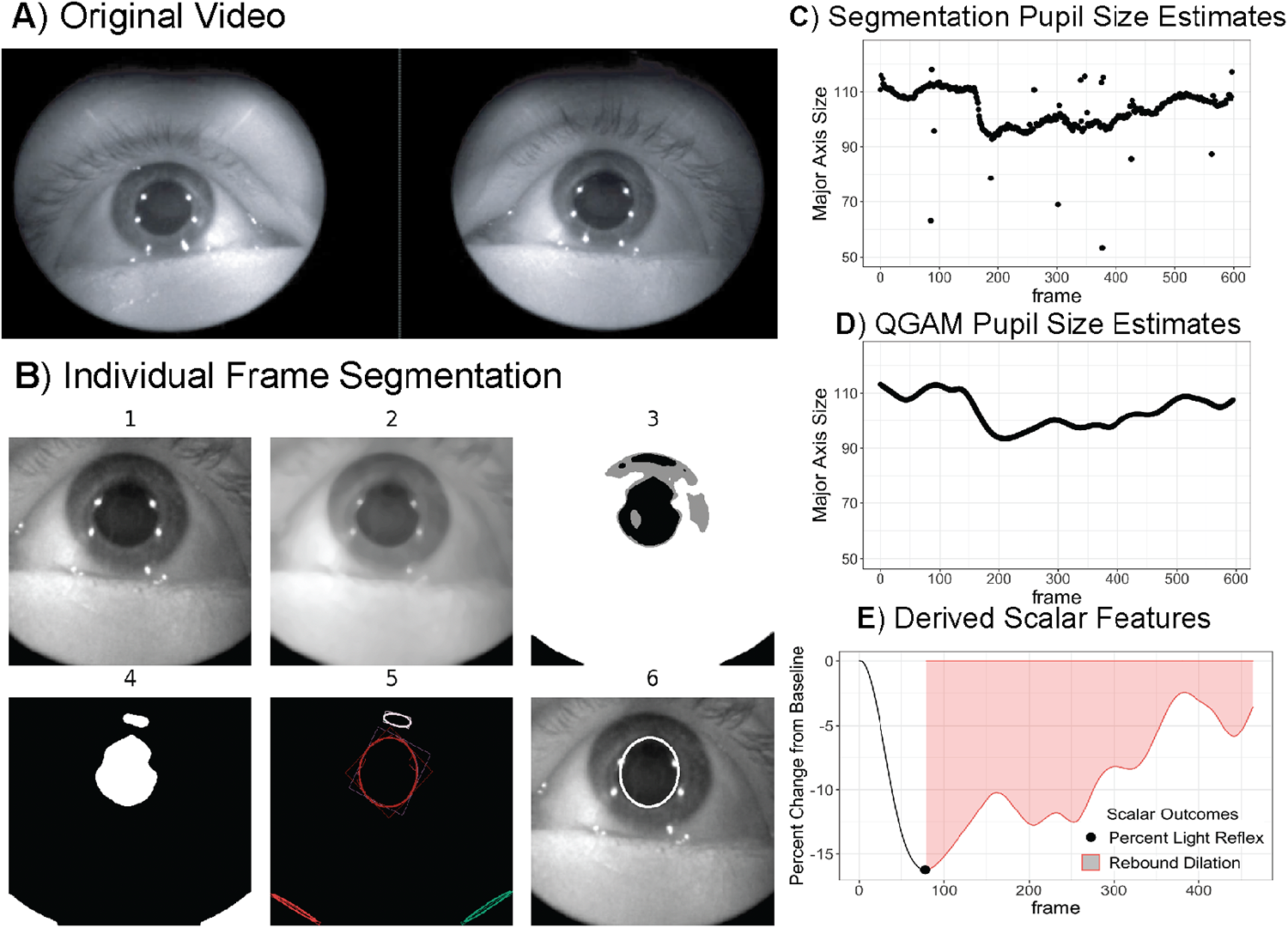
**A** shows a full frame captured from a video at the beginning of the light stimulus test. **B** displays an abbreviated version of the segmentation pipeline used to predict pupil size for each eye in each frame. In **C**, pupil size estimates from the segmentation pipeline are plotted for one example video where major axis size is measured in pixels. In **D**, pupil size estimates from the quantile additive regression model are plotted displaying a smoothed trajectory using the data from **C**. In **E**, two scalar features that we analyze are displayed: percent light reflex and rebound dilation.

**Figure 2:**
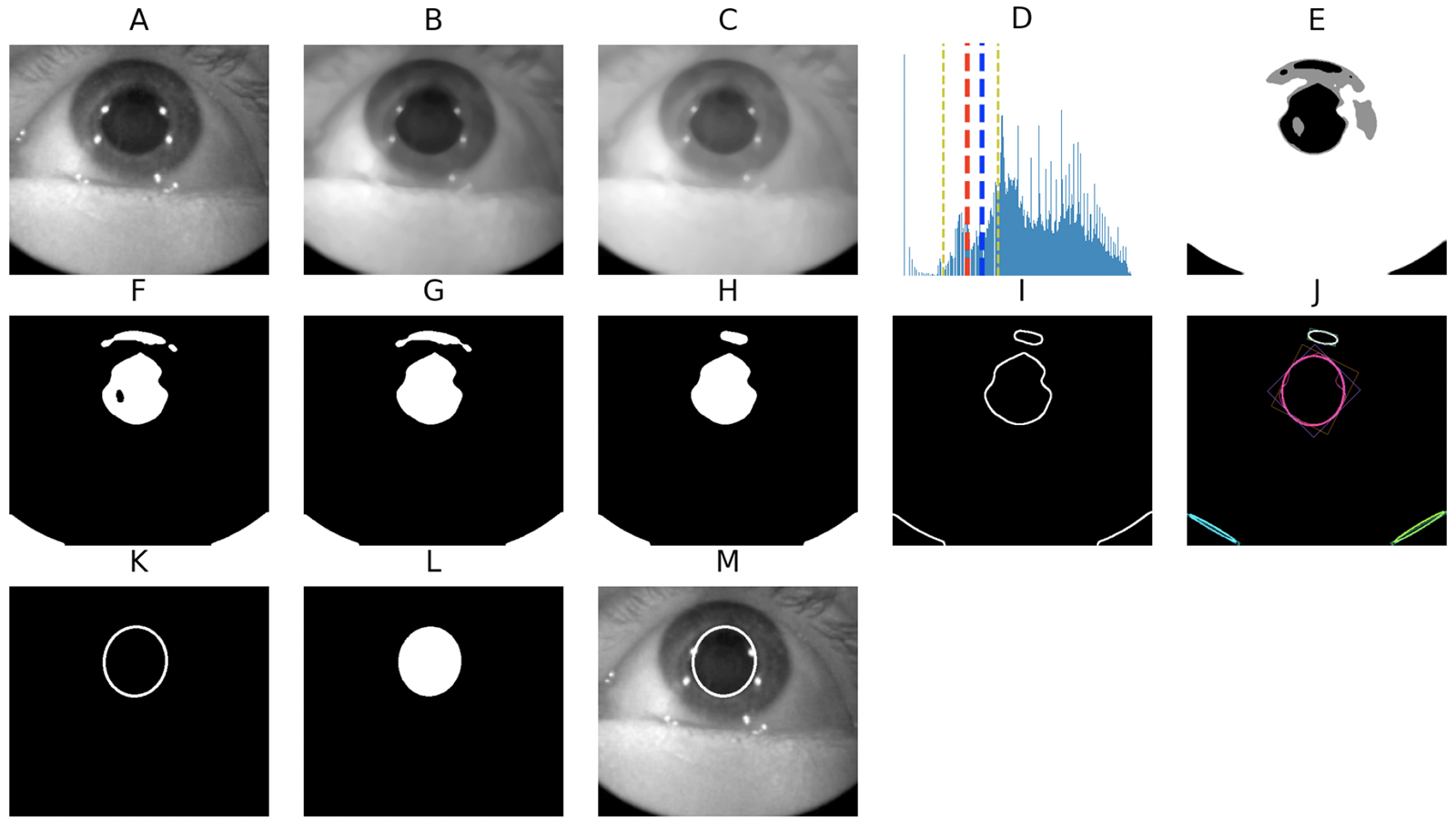
The process for creating a binary pupil mask from an image. Panel **A** shows original image. **B** and **C** show the image after applying a smoothing median kernel (size 15) and a power law intensity transform (*γ* = 0.7), respectively. **D** shows the histogram of pixel intensities for the image shown in **C**. The two dashed yellow lines represent a truncated subset of the pixel intensities where we search for intensity thresholds that separate the pupil, iris, and background. **E** shows the image after applying thresholds obtained in **D**. In **F**, a binary image is created by selecting the lowest intensity class from **E. G** and **H** show the binary image after it has undergone an image morphology closing and opening, respectively. **I** shows the image after edge detection. In **J**, a minimum ellipse was fit around each contour. **K** shows the “best ellipse” selected based on area, eccentricity, and location. **L** shows the final binary mask. Finally, **M** shows the segmentation results from **L** projected onto the original image.

First a median smoothing filter was convolved over each image to smooth noise that arises from intensity differences across videos (Figure 2**B**). Next, a power law transformation defined by 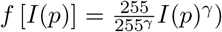 was applied to each image with intensity transform constant *γ* (Figure 2**C**). The power law intensity transformation modifies pixel intensity values to increase contrast in lighter (*γ >* 1) or darker (*γ* < 1) regions of the image. Next the image was segmented using Otsu’s multi-level thresholding algorithm. Otsu’s algorithm is applied to an image intensity histogram and distinguishes *k*-classes by maximizing inter-class variance (Otsu, 1979). For Otsu’s method, we truncated the intensity histogram to only consider pixels from the 1st to 20th quantile which created more balanced classes and enhanced segmentation performance (Figure **2D**). Our *k* = 3 classes separated images into intensity distributions with the lowest, middle, and highest intensity values roughly describing the pupil, iris, and background pixels (Figure **2E**). Next we created a binary mask by designating the lowest intensity class from the Otsu method as the pupil pixels (Figure **2F**), and performed an image morphology closing and opening operation (Figures **2G** and **2H**) to close any holes and separate any noise arising from the iris or eyelashes (Haralick et al., 1987).

Canny edge detection, a method for finding image structure using gradient filters, was then applied to the processed binary image in Figure **2I** (Ding and Goshtasby, 2001). From the edges in each image we fit ellipses (Figure **2J**) using the LIN algorithm for algebraic distance, which minimizes the squared distance between the observed edge points and fitted ellipse (Fitzgibbon et al., 1996). Each image could contain multiple ellipses, and we calculated the center, eccentricity, and the major and minor axis for each ellipse in each image. The ellipses were filtered by eccentricity to ensure estimated pupils are relatively circular, and by area to ensure biologically plausible size (Figure **2K**). We used the manually segmented data to define realistic ranges for eccentricity and area; specifically, **ellipse eccentricity** 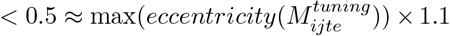 and 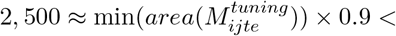 **ellipse area** 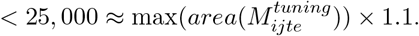.

Since it was possible for multiple ellipses to fit these criteria within a single image, the ellipse that was closest to the center of the frame was accepted as the best estimate for the pupil and used to compute the binary mask, 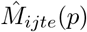 (Figure **2M**.)

For a small quantity of images, the iris pixel intensities overlapped with the pupil pixel intensities, resulting in the pupil being connected to pieces of the iris even after image morphology (Supplement Figure 3). If the pipeline described above failed to fit an ellipse satisfying area, eccentricity, and location requirements, the same pipeline was run with an additional step. After image morphology, the Watershed region-based segmentation algorithm (Beucher and Meyer, 2018), was applied to separate the binary image into multiple component pieces (Supplement Figure 3**C**). For each component piece, a minimum ellipse was fit, and a final pupil size prediction was made using the same ellipse filtering process described above. (Supplement Figures 3**D**-3**H**). If no ellipse was successfully fit, pupil size was recorded as “NA”. The failure to fit an ellipse most commonly occurred in frames where the subject was blinking.

The tuning set of manually segmented masks, 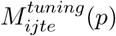, was used to determine the following segmentation parameters: median kernel size, gamma transformation value, quantiles at which to truncate the intensity histogram for Otsu thresholding, disk size for image morphology, and watershed minimum basin size. A grid search over potential parameter values was performed and optimal values were chosen that maximized the mean F-score over all images in the tuning set. The segmentation pipeline using the selected parameters was then applied to each frame across all videos. The major axis size from each final ellipse was used in subsequent analysis.

#### 2.2.3 Processing Pupil Size Trajectories

In the segmentation step we estimated pupil major axis size in pixels from each frame of each video. This results in a pupil size trajectory for each eye from every video, which describes the behavior of the pupil over the duration of the light stimulus test. This section describes how pupil size trajectories, created were cleaned and smoothed using quantile additive regression models.

Each pupil size trajectory contained some extreme outliers which were not due to biological variability in pupil size, but instead, were due to artifacts like blinking or pixel intensity overlap between the pupil and iris (Figure 3**A**). To remove these outliers from our pupil size trajectories while still capturing biological nonlinear fluctuations in pupil size, we fit a smooth additive median regression model (QGAM) to each trajectory (Fasiolo et al., 2021a). Any pupil major axis sizes further than 6 standard deviations away from the QGAM predicted values were removed as outliers (Figure **3B**). We then refit the additive median regression model on the cleaned data, and used predicted values from this model to derive scalar features. (Figure **3C**).

**Figure 3:**
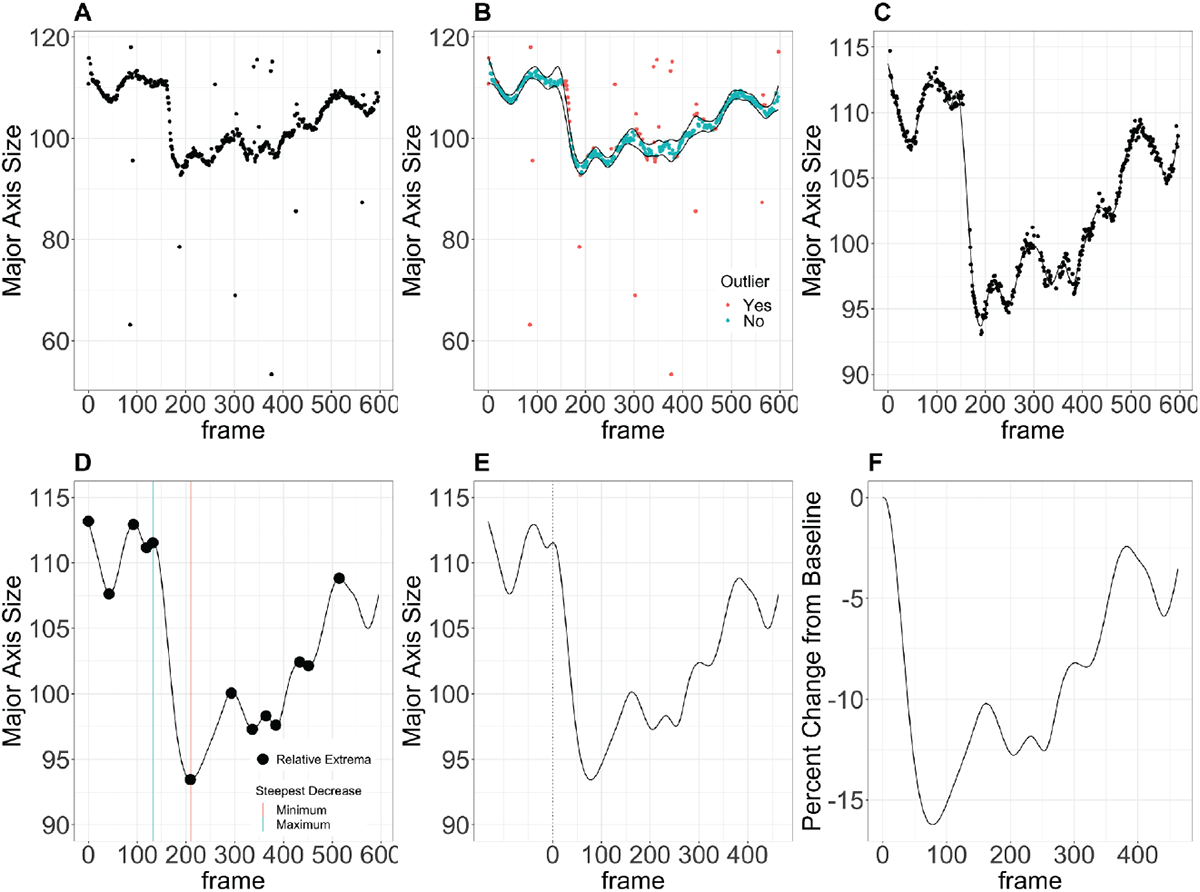
**A** shows the estimated major axis size for the right eye from one video. **B** shows the estimated major axis size ± 6 standard deviations from the median of a QGAM fit and highlights points that were identified as outliers. **C** shows the cleaned pupil trajectory (points) and QGAM fit (line) **B. D** shows the relative extremum along the pupil trajectory and two lines indicating the set of relative extremum that captures the steepest decrease in the first 300 frames of the video. **E** shows the shifted frames to set the maximum point from **D** as frame zero. **F** shows the final registered curve, which shows only frames from the start to the end of the light response test.

To evaluate the success of the cleaning process we calculated mean autocorrelation for each pupil trajectory before and after outlier removal. High autocorrelation implies neighboring pupil size estimates were similar, which suggests smooth trajectories for pupil size over time. Out of 202 videos, trajectories from 47 videos were identified as requiring manual cleaning because they had an autocorrelation below 0.7 after outlier removal. Trajectories from these 47 videos were discussed until the research team came to consensus about whether to keep or remove the curves. The manual cleaning process entailed looking at the pupil size trajectories and determining which pupil size estimates were likely not part of the true biological response. Estimates for pupil size that were deemed not part of the true pupil size trajectory were removed by assigning them as an outlier and filtering the data to include only non-outliers using R. Forty of the 47 videos had a pupil size trajectory for at least one eye that could be recognized and cleaned. Seven videos were removed from analysis that did not have clear trajectories for either eye.

On average, the cleaning process removed 146 out of 600 observations from each trajectory, and increased mean autocorrelation for each video from 0.804 to 0.966.

#### 2.2.4 Pupil Trajectory Alignment

To conduct statistical inference on the pupil size trajectories, the curves first had to be aligned, or registered, to the beginning of the light stimulus test. This required estimating the frame at which the light response test began and ended for each video. To detect the start of the test we first located all relative extrema in the first 300 frames of the pupil trajectory and found the largest decrease between extrema (Figure **3D**). The maximum before the largest decrease was then designated the start of the light test and appointed frame zero (Figure **3E**). To detect the end of the light test, we leveraged the information that participants were instructed to close their eyes at the end of the test. At that moment, estimates of pupil size see a large fluctuation from their more stable trajectory. We end the test for each subject at the frame where the preceding frame was within five pixels and the two successive frames were, on average, ten pixels or more away. This ending ensured the final estimate was close to the estimated trajectory and removes points that significantly differed from the estimated trajectory. Percent change from the start of the test was then calculated for each frame (Figure 3**F**).

#### 2.2.5 Extraction of Pupil Response Features

Scalar features were calculated from each pupil size trajectory to compare pupil response at the pre and post assessments and across cannabis use groups. Percent light reflex was defined as the magnitude of peak decrease in pupil diameter during illumination as a percentage of the pre-illumination diameter. Thus, a high “percent reflex” denotes greater illumination induced pupillary constriction. After a pupil achieves its maximal constriction (smallest diameter) during illumination, it may in some cases subsequently dilate and increase in diameter during ongoing illumination. This process is termed “rebound dilation.” The magnitude of rebound dilation in this study was assessed by calculating the area under the curve (AUC) of percent change in pupil size after the point of minimum constriction. Because illumination time was found to vary (IQR: 12.7, 15.0 seconds) across videos, the calculated AUC was standardized by dividing by the total number of frames considered. Rebound dilation is thus reported as percent pupil diameter change from baseline per frame, with negative values indicating pupillary constriction. The more negative a value, the larger the area under the curve for pupil constriction over time of illumination, and hence the less “rebound dilation” that occurred.

#### 2.2.6 Statistical Analyses

Segmentation performance was evaluated by comparing the estimated binary masks, 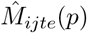, with the validation set of manually segmented masks, 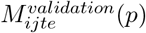. We used accuracy, precision, Jaccard index, and F-score metrics to analyze the validity of our segmentation pipeline. True positives and true negatives were defined as correctly identifying a pixel as pupil or background, respectively. Scalar features were modeled using generalized estimating equations with a Gaussian link function and independent correlation structure. The predictors for each model included assessment period (pre or post), cannabis use group, eye, and an interaction between cannabis use group and assessment period. Wald p-values are reported, and contrasts were used to statistically test differences in pre to post assessment outcome values between groups.

All video processing and segmentation analyses were conducted using Python version 3.8.8. The `Scikit-image` module was used for image processing steps including image filtering and morphology (van der Walt et al., 2014), and the `OpenCV` module was used for video decomposition and ellipse fitting (Bradski, 2000). Statistical analyses were conducting using R Studio version 1.3.1073. Additive median regression models were fit using cubic splines with 25 knows using the `qgam` package (Fasiolo et al., 2021b), and the `geepack` package was used to fit generalized estimating equations (Halekoh et al., 2006). Analyses were implemented on a standard laptop. The code for this segmentation pipeline is available on GitHub.

## 3 Results

### 3.1 Parameter Selection and Computation Time

Parameters for the final segmentation model were selected using a grid search to maximize F-score in the tuning images. Using this approach we selected a size 15 kernel for the median filter, *γ* = 0.7 for the power-law intensity transform, and we truncated the intensity histogram to only include values between the 1st and 20th quantiles for Otsu’s multi-level thresholding. Disk size for image morphology and watershed minimum basin size were selected to be 8 and 25, respectively. These parameter values were implemented in the segmentation pipeline and applied to the full data set of 202 videos, 7 of which were removed from the data set after segmentation and manual cleaning due to quality control issues. In some cases data was only usable for one of the two eyes; as a result we retained pupil trajectories for both eyes from 179 videos, from the left eye only for 9 videos, and from the right eye only for 6 videos. Computation time was, on average, 6 minutes and 15 seconds per video.

### 3.2 Pupil Segmentation

Segmentation performance as measured by accuracy, precision, F-score, and Jaccard index on both the tuning and validating sets is presented in Table 1. On successfully segmented frames, our segmentation model performed well across all metrics, with particularly high accuracy (validating set median [IQR] of 0.995 [0.992, 0.997]) and precision (validating set median [IQR] of 0.997 [0.985, 1]). F-score was used to optimize our pipeline parameters and achieved a median [IQR] of 0.944 [0.922, 0.96] on the validating set. The Jaccard index (validating set median [IQR] of 0.894 [0.855, 0.923]) was slightly lower than other metrics. However, a high Jaccard index was still achieved despite an inherent bias in the minimum ellipse fitting algorithm to under fit the points slightly, leading to an upper limit for the Jaccard index less than 1. This bias to under fit is not concerning as it is consistent across all images. As seen in Table 1, the segmentation pipeline performed consistently across the tuning and validating data sets. Together, these results suggest stable, successful segmentation.

**Table 1:** Segmentation Results. Segmentation results comparing binary pupil masks estimated from our model with manually segmented masks. On the tuning set consisting of 1217 images, 1213 ellipses were successfully fit. On the validating set, 517 ellipses were successfully fit out of 517 images. Results are consistent on the tuning and validating sets. On the validating set, the median of accuracy, precision, F-Score, and Jaccard index were 0.995, 0.997, 0.944, and 0.894, respectively.

| Metric | Tuning (N=1213) | Validating (N = 517) | Overall (N = 1730) |
| --- | --- | --- | --- |
| Accuracy | 0.995 (0.992, 0.998) | 0.995 (0.992, 0.997) | 0.995 (0.992, 0.997) |
| Precision | 0.992 (0.977, 1) | 0.997 (0.985, 1) | 0.994 (0.98, 1) |
| F-Score | 0.951 (0.926, 0.967) | 0.944 (0.922, 0.96) | 0.949 (0.924, 0.966) |
| Jaccard index | 0.907 (0.862, 0.937) | 0.894 (0.855, 0.923) | 0.903 (0.858, 0.933) |

### 3.3 Analysis of Pupil Response Features

Eye laterality did not affect the model for percent light reflex (p = 0.687) or rebound dilation (p = 0.304), indicating that neither scalar feature differed substantially between right and left eyes; however, results reported in Tables 2 and 3 are adjusted for eye.

**Table 2:** Scalar Feature Summary. Mean (standard deviation) of percent light reflex and mean (standard deviation) of rebound dilation values for the cannabis no use, occasional use, and daily use groups at baseline (“pre”) and post-smoking (“post”). Differences across use groups were not statistically significant at the pre or post measuring event.

| Use Group | *Baseline* |  | *Post-Smoking* |  |
| --- | --- | --- | --- | --- |
|  | Percent Light Reflex | Rebound Dilation | Percent Light Reflex | Rebound Dilation |
| No Use | 29.6 (1.53) | -14.8 (1.28) | 23.7 (2.77) | -11.3 (2.65) |
| Occasional | 33.8 (3.72) | -17.9 (3) | 20.4 (6.93) | -11.1 (6.2) |
| Daily | 32.9 (3.85) | -19.3 (3.28) | 20.1 (7.09) | -11.3 (6.57) |
| Any | 33.4 (3.46) | -18.6 (2.87) | 20.2 (6.35) | -11.2 (5.88) |

**Table 3:**
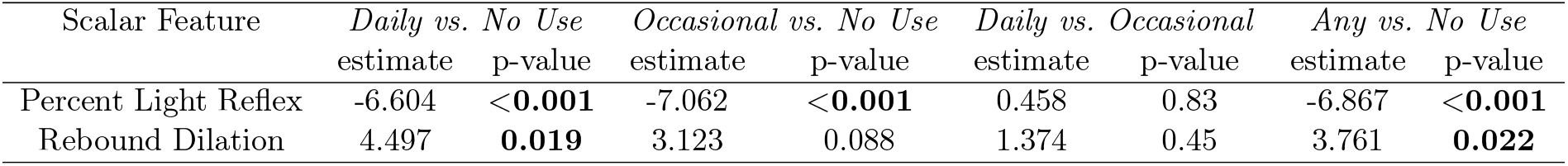
Use Group Comparison Summary. Estimated effects and p-values for pairwise comparisons between cannabis use groups. The estimates reflect the difference in change from pre to post, where the second group listed was the reference group, and the difference was calculated as comparison change minus reference change.

| Scalar Feature | *Daily vs. No Use* |  | *Occasional vs. No Use* |  | *Daily vs. Occasional* |  | *Any vs. No Use* |  |
| --- | --- | --- | --- | --- | --- | --- | --- | --- |
|  | estimate | p-value | estimate | p-value | estimate | p-value | estimate | p-value |
| Percent Light Reflex | -6.604 | **<0.001** | -7.062 | **<0.001** | 0.458 | 0.83 | -6.867 | **<0.001** |
| Rebound Dilation | 4.497 | **0.019** | 3.123 | 0.088 | 1.374 | 0.45 | 3.761 | **0.022** |

Table 2 presents adjusted means and standard errors for percent light reflex and for comparison across cannabis use groups at baseline and after smoking. The “any use” group presents the average between daily and occasional use groups. At the baseline assessment, mean (SD) percent light reflex for no, occasional, and daily use groups was 29.6 (1.53), 33.8 (3.72), and 32.9 (3.85), respectively. Those with no use exhibited 23.7 (2.77) percent light reflex at the “post” assessment, somewhat greater than that observed at the post-smoking time point in occasional and daily use groups, whose percent light reflex were 20.4 (6.93) and 20.1 (7.09) percent, respectively. However, percent light reflex did not statistically differ by user group at either the baseline or post-smoking time points (*p* = 0.25, *p* = 0.3, respectively).

At the baseline assessment rebound dilation values were -14.8 (1.28), -17.9 (3), and -19.3 (3.28) for the no, occasional, and daily use groups respectively. Rebound dilation at the post-smoking assessment was similar across groups with values of -11.3 (2.65), -11.1 (6.2), -11.3 (6.57), respectively. Rebound dilation did not statistically differ by user group at either baseline or post-smoking time points (*p* = 0.09, *p* = 1, respectively). High standard deviations for both percent light reflex and rebound dilation indicate heterogeneity among individuals that may mask differences across cannabis use groups; for this reason, we focus more on within subject differences presented below.

Table 3 contains the adjusted mean difference of differences (post smoking - baseline) in outcome variables for each pairwise combination of use group. The mean change in percent light reflex was significantly larger for those with occasional use relative to no use (*p* < 0.001) and daily use relative to no use (*p* < 0.001), indicating less pupil constriction after smoking for those with daily and occasional use than for no use. There was not a significant change in mean percent light reflex comparing those with occasional use to daily use (*p* = 0.83). The mean change in rebound dilation was significantly larger for those with daily use relative to no use (*p* = 0.019), but not for those with occasional use relative to no use (*p* = 0.088), or occasional use relative to daily use (*p* = 0.45). Collapsing daily and occasional use into one “any use” group, both the percent light reflex (*p* < 0.001) and rebound dilation (*p* = 0.022) outcomes were significantly different for any use compared to no use.

## 4 Discussion

We present a complete pipeline for the segmentation of pupils and analysis of cannabis-associated changes in pupillary size using data captured from infrared video of pupils exposed to visible light illumination. Using objective measurements, we found differences in pupil response after cannabis use for those with daily and occasional use.

Our results indicate that acute cannabis smoking reduced the percent light reflex in both occasional and daily smokers, a finding consistent with prior reports (Fant et al., 1998; Campobasso et al., 2020). We also found that rebound dilation, the physiological re-dilation that occurs despite sustained eye illumination, increased after acute cannabis smoking. These changes in pupillary response to illumination did not differ significantly when those with occasional use were compared to those with daily use. The lack of difference between these two types of cannabis use groups suggests that tolerance to cannabis-induced changes in pupillary light response may not occur. This is in contrast to tolerance-associated differences between those with a history of occasional versus daily use for a variety of cognitive or cardiac effects (Colizzi and Bhattacharyya, 2018).

This study was subject to several limitations. The software included with the apparatus did not provide illumination for a consistent interval of time between tests, and did not indicate at what time frame the light response test began and ended. As such, these quantities had to be estimated within our pipeline, which introduced additional uncertainty. Another limitation was that the camera-to-eye distance associated with the stand-mounted goggles used for this study was unmeasured, thus preventing quantitative estimation of pupil diameter in millimeters. Prior research has demonstrated that baseline pupil diameter is an independent predictor of the magnitude of pupillary light reflex, in that greater percent constriction occurs post-illumination in pupils as baseline pupil diameter increases (McKay et al., 2018; Larson and Behrends, 2015). Accordingly, without any ability to include baseline pupil diameter as an independent variable in the regression analysis, we are unable to determine how much of the relative change in percent light reflex between user groups was due to an acute effect of cannabis as opposed to quantitative differences in baseline (pre-illumination) pupil diameter.

Prior research on the impact of acute cannabis smoking on pupil size, aside from reporting qualitative inconsistencies (miosis, mydriasis or no change), has indicated that cannabis-associated change in pupil size, when it does occur, is rather small, i.e., < 0.3 mm (Ortiz-Peregrina et al., 2020). From a practical standpoint, changes of this size would be difficult to accurately or precisely measure without the use of sensitive quantitative infrared pupillometers. Moreover, because the sparse published data available suggests that the variance observed in pupil diameter in normal healthy subjects in the drug-free state may equal or exceed the magnitude of acute change associated with cannabis smoking, it may be challenging to utilize single measurements of pupil size or pupillary light reflex to accurately assess if subjects were likely to have recently smoked cannabis. In the present study, which lacked determination of pupil diameter in millimeters, there was no significant difference in percent light reflex or rebound dilation at the post smoking time point (or second measurement) between control subjects and those who had just smoked cannabis. Additional research conducted with sensitive quantitative infrared pupillometry that contributes to larger normative data sets and measures time dependent change in pupillary parameters associated with acute cannabis consumption will be helpful to assess the ultimate utility of pupillometry as a biomarker of cannabis use.

## A Appendix section

The datasets and code used for this project can be found at https://github.com/yf297/GpGp_*m*_*ulti*_*p*_*aper*.∗ *updatethis*

## Acknowledgments

We would like to thank James Sherrick for completing the manual segmentation of images.

## References

Beucher S, Meyer F (2018). The morphological approach to segmentation: the watershed transformation. In: Mathematical morphology in image processing, 433–481. CRC Press.

Bradski G (2000). The OpenCV Library. Dr. Dobb’s Journal of Software Tools.

Brooks-Russell A, Brown T, Friedman K, Wrobel J, Schwarz J, Dooley G, et al. (2021). Simulated driving performance among daily and occasional cannabis users. Accident Analysis & Prevention, 160: 106326.

Brown B, Adams AJ, Haegerstrom-Portnoy G, Jones RT, Flom MC (1977). Pupil size after use of marijuana and alcohol. American journal of ophthalmology, 83(3): 350–354.

Brubacher JR, Chan H, Erdelyi S, Macdonald S, Asbridge M, Mann RE, et al. (2019). Cannabis use as a risk factor for causing motor vehicle crashes: a prospective study. Addiction, 114(9): 1616–1626.

Campobasso CP, De Micco F, Corbi G, Keller T, Hartung B, Daldrup T, et al. (2020). Pupillary effects in habitual cannabis consumers quantified with pupillography. Forensic Science International, 317: 110559.

Colizzi M, Bhattacharyya S (2018). Cannabis use and the development of tolerance: a systematic review of human evidence. Neuroscience & Biobehavioral Reviews, 93: 1–25.

Ding L, Goshtasby A (2001). On the canny edge detector. Pattern recognition, 34(3): 721–725.

Downey LA, King R, Papafotiou K, Swann P, Ogden E, Boorman M, et al. (2012). Detecting impairment associated with cannabis with and without alcohol on the standardized field sobriety tests. Psychopharmacology, 224(4): 581–589.

Fant RV, Heishman SJ, Bunker EB, Pickworth WB (1998). Acute and residual effects of marijuana in humans. Pharmacology Biochemistry and Behavior, 60(4): 777–784.

Fasiolo M, Wood SN, Zaffran M, Nedellec R, Goude Y (2021a). Fast calibrated additive quantile regression. Journal of the American Statistical Association, 116(535): 1402–1412.

Fasiolo M, Wood SN, Zaffran M, Nedellec R, Goude Y (2021b). qgam: Bayesian nonparametric quantile regression modeling in R. Journal of Statistical Software, 100(9): 1–31.

Fitzgibbon AW, Fisher RB, et al. (1996). A buyer’s guide to conic fitting. Citeseer.

Halekoh U, Højsgaard S, Yan J (2006). The r package geepack for generalized estimating equations. Journal of Statistical Software, 15/2: 1–11.

Haralick RM, Sternberg SR, Zhuang X (1987). Image analysis using mathematical morphology. IEEE transactions on pattern analysis and machine intelligence, (4): 532–550.

Hartman RL, Richman JE, Hayes CE, Huestis MA (2016). Drug recognition expert (dre) examination characteristics of cannabis impairment. Accident Analysis & Prevention, 92: 219–229. IACP (2022). The 12-step dre protocol. https://www.theiacp.org/12-step-process. Accessed: 2022-05-30.

Lamprecht MR, Sabatini DM, Carpenter AE (2007). Cellprofiler™: free, versatile software for automated biological image analysis. Biotechniques, 42(1): 71–75.

Larson MD, Behrends M (2015). Portable infrared pupillometry: a review. Anesthesia & Analgesia, 120(6): 1242–1253.

Lira MC, Heeren TC, Buczek M, Blanchette JG, Smart R, Pacula RL, et al. (2021). Trends in cannabis involvement and risk of alcohol involvement in motor vehicle crash fatalities in the united states, 2000–2018. American journal of public health, 111(11): 1976–1985.

Logan B, Kacinko SL, Beirness DJ (2016). An evaluation of data from drivers arrested for driving under the influence in relation to per se limits for cannabis.

McKay RE, Neice AE, Larson MD (2018). Pupillary unrest in ambient light and prediction of opioid responsiveness: case report on its utility in the management of 2 patients with challenging acute pain conditions. A&A Practice, 10(10): 279–282.

Merzouki A, Mesa JM, Louktibi A, Kadiri M, Urbano G (2008). Assessing changes in pupillary size in rifian smokers of kif (cannabis sativa l.). Journal of forensic and Legal Medicine, 15(5): 335–338.

Newmeyer MN, Swortwood MJ, Taylor ME, Abulseoud OA, Woodward TH, Huestis MA (2017). Evaluation of divided attention psychophysical task performance and effects on pupil sizes following smoked, vaporized and oral cannabis administration. Journal of applied toxicology, 37(8): 922–932.

Ortiz-Peregrina S, Ortiz C, Castro-Torres JJ, Jiménez JR, Anera RG (2020). Effects of smoking cannabis on visual function and driving performance. a driving-simulator based study. International journal of environmental research and public health, 17(23): 9033.

Otsu N (1979). A threshold selection method from gray-level histograms. IEEE transactions on systems, man, and cybernetics, 9(1): 62–66.

Papafotiou K, Carter JD, Stough C (2005). An evaluation of the sensitivity of the standardised field sobriety tests (sfsts) to detect impairment due to marijuana intoxication. Psychopharmacology, 180(1): 107–114.

Shahidi Zandi A, Comeau FJ, Mann RE, Di Ciano P, Arslan EP, Murphy T, et al. (2021). Preliminary eye-tracking data as a nonintrusive marker for blood δ-9-tetrahydrocannabinol concentration and drugged driving. Cannabis and cannabinoid research, 6(6): 537–547.

Stark M, Englehart K, Sexton B, Tunbridge R, Jackson P (2003). Use of a pupillometer to assess change in pupillary size post-cannabis. Journal of Clinical Forensic Medicine, 10(1): 9–11.

van der Walt S, Schönberger JL, Nunez-Iglesias J, Boulogne F, Warner JD, Yager N, et al. (2014). scikit-image: image processing in Python. PeerJ, 2: e453.

